# Screening of a kinase inhibitor library in human Huntington’s disease iPSC-derived striatal precursor neurons reveals a neuroprotective effect of PKC-α/β1 inhibition

**DOI:** 10.1101/2025.09.18.677178

**Authors:** Mali Jiang, Tianze Shi, Ritika Miryala, Matthew Rodriguez, Ronald Wang, Rashi Sultania, Lauren Guttman, Julia Johnston, Louisa Mulligan, Yushan Li, Anning Cui, Karim Belkas, Yuxuan Xue, Yuna Um, Ammy Yuan, Chloe Holland, Juan C. Troncoso, Wenzhen Duan, Tamara Ratovitski, Wanli Smith, Christopher A. Ross

**Author notes:** These authors contributed equally (Mali Jiang, Tianze Shi, Ritika Miryala). In brief: The loss of striatal neurons is a defining feature of Huntington’s disease (HD). Using a human striatal neuronal model derived from HD patient iPSCs, we identified the PKC-α/β1 inhibitor GO6976 as a protective agent against mutant huntingtin toxicity, positioning PKC-α/β1 as a potential therapeutic target for HD. Correspondence (M.J.) (C.R.).

## Abstract

The loss of striatal medium spiny neurons is a hallmark of Huntington’s disease (HD). To identify potential disease-modifying treatments, we previously developed a human neuronal model by immortalizing and differentiating HD patient-derived iPSCs into highly homogeneous striatal precursor neurons (ISPNs). Using a 96-well screening platform, and two rounds of re-screening, we tested a kinase inhibitor library and identified 5 compounds that protected HD ISPNs from mutant huntingtin (mHTT)-induced toxicity. Among these, we prioritized the PKC-α/β1 inhibitor GO6976, which rescued HD ISPNs from mHTT toxicity in a dose-dependent manner. Further, we found increased phosphorylation of PKC-α and PKC-β1 in HD cells and tissues, while their overexpression was toxic to HD ISPNs. Knockdown of PKC-α/β1 protected the neurons, and both isoforms interacted and colocalized with HTT. These results suggest that PKC-α/β1 plays a role in HD neurodegeneration, and that inhibiting their activity may offer a potential therapeutic approach for HD.

**Graphical abstract:** 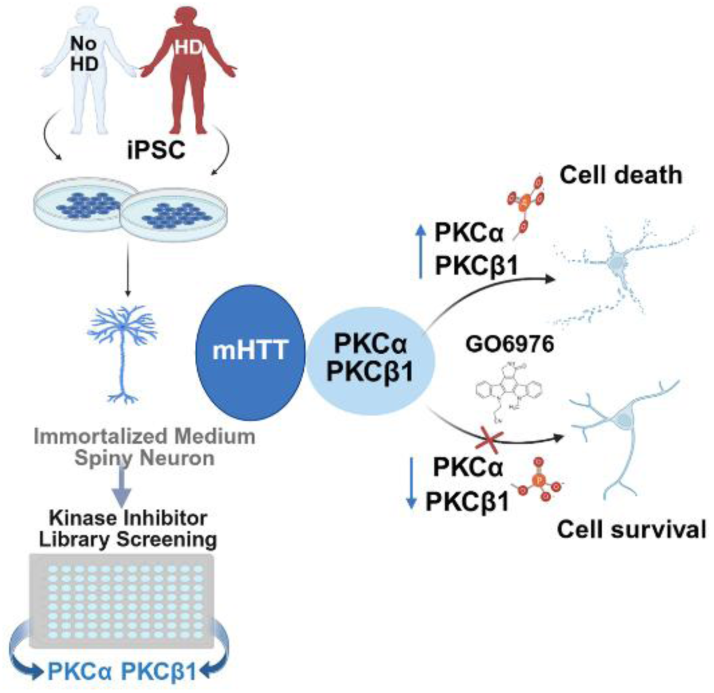

**Highlights:**

- HD patient-derived iPSC-based striatal precursor neurons (ISPNs) were used to screen and identify neuroprotective compounds.
- The PKC-α/β1 inhibitor GO6976 rescues HD ISPNs from mutant huntingtin (HTT)-induced toxicity.
- The phosphorylation of PKC-α/β1 is elevated in HD cell and tissues, and PKC-α/β1 interact with both wild-type and mutant huntingtin.
- Overexpression of PKC-α/β1 is toxic to HD ISPNs, while its knockdown protects the neurons.

## Introduction

Huntington’s disease (HD) is an autosomal dominant neurodegenerative disorder caused by an expanded CAG repeat (> 36) in exon 1 of the *Huntingtin* (*HTT)* gene, resulting in an extended polyglutamine tract in the huntingtin protein (HTT) ^1–4^. Clinically, HD is characterized by involuntary choreiform movements, voluntary movement dysfunction, psychiatric changes, and cognitive impairment ^5–7^. Although the HTT protein has diverse functions and is ubiquitously expressed, the hallmark neuropathology of HD is the loss of medium spiny neurons (MSNs) in the striatum ^8–10^. Currently, there is no disease modifying therapy available for HD.

Most recent therapeutic strategies for HD have focused on lowering HTT levels and inhibiting DNA mismatch repair (MMR)^6^. However, the inability to distinguish between wild-type and mutant HTT raises safety concerns, and the lack of efficacy observed in clinical trials has set back the HTT lowering approach ^11,12^. Small-molecule MMR inhibitors offer a promising strategy to reduce somatic instability (which is an initial step in mutant *HTT* pathogenesis), but their effectiveness may be limited to the early stages of the disease ^13,14^. Therefore, additional targets for disease modifying neuroprotective agents are urgently needed—either as standalone therapies or to complement other approaches.

HD cell models represent useful tools for understanding the molecular mechanism of the disorder and evaluating potential therapeutics. Among these, human iPSCs offer significant advantages over traditional cell models, particularly in disease modeling, drug screening, and cell replacement therapy ^15–17^. Emerging evidence suggests that CAG repeat expansions and their associated toxicity exhibit cell-type selectivity in MSNs of HD patients, contributing to neurodegeneration ^18,19^. HD iPSCs generated from HD patients with different CAG repeat lengths can be differentiated to an MSN-like phenotype. This system recapitulates aspects of HD-related cellular pathogenesis ^20–22^. However, several limitations of HD iPSCs hinder their broader application, including long and complicated differentiation protocols, limited yield of differentiated cells, and potential variability in cell phenotypes among different experiments. To overcome these disadvantages, we previously established a human neuronal model by immortalizing and differentiating HD patient iPSCs into highly homogeneous striatal precursor neurons (ISPN), which can be further differentiated to an MSN-like phenotype over a two-week period. This model recapitulates medium spiny phenotypes observed in parental iPSCs, including MAP2 and DARPP32 expression ^23^.

Recent work by the McCarroll and Heintz labs integrated CAG repeat sizing with RNA-seq at the single-cell level and across different cell types, revealing that somatic CAG expansion plays a central role in HD pathogenesis ^18,19^. The McCarrol’s findings indicate that MSNs with up to 150 CAG repeats show minimal gene expression changes, while those with over 150 repeats exhibit loss of key MSN-defining transcripts and activation of aberrant developmental or toxic pathways ^19^. Our ISPN HD model, featuring over 180 CAG repeats, is uniquely positioned to study mHTT toxicity after somatic expansion. Thus, ISPNs derived from HD iPSCs provide a robust and reproducible cell system for drug screening and target validation, offering a valuable resource in the fight against HD.

To identify small molecules as potential disease-modifying treatments for HD using human cell-based assays, we developed a 96-well plate screening platform utilizing the CellTiter-Glo luminescent cell viability assay in ISPNs. A kinase inhibitor library (ApexBio) containing 764 compounds was screened in HD ISPNs expressing 180 CAG repeats (180Q-ISPNs). This screening identified 5 compounds that protected HD ISPNs from mHTT-induced toxicity in the context of cellular stress. Among the hits, we prioritized GO6976, a small molecule inhibitor of PKC-α and PKC-β1. Further validation confirmed that PKC-α and PKC-β1 play significant roles in HD pathogenesis.

## Results

### Drug screening platform utilizing HD striatal precursor neurons derived from human iPSCs

We previously established human striatal ISPN lines derived from HD patient iPSCs, which recapitulate key HD-related phenotypes of the parental iPSC model. Building on this, we developed an assay utilizing the CellTiter-Glo luminescent cell viability reagents in mature striatal neurons differentiated from ISPNs (Figure 1A). Our findings indicate that striatal neurons with 180 CAG repeats exhibit greater vulnerability to BDNF withdrawal-induced stress compared to neurons with 33 CAG repeats. Notably, HD ISPN striatal precursor neurons show similar sensitivity to growth factor and nutrient withdrawal as their mature counterparts (Figure 1B). Using this system, we established a mutant HTT cellular toxicity 96-well screening platform for target validation and drug screening (Figure 1C). The assays achieved Z’ scores > 0.5, validating the robustness of the platform. The vulnerability of HD ISPNs was further validated using the Caspase-Glo 3/7 assay (Figure S1). Nevertheless, owing to considerable inter-experimental variability, this assay was deemed not adopted for subsequent screening.

**Figure 1.**
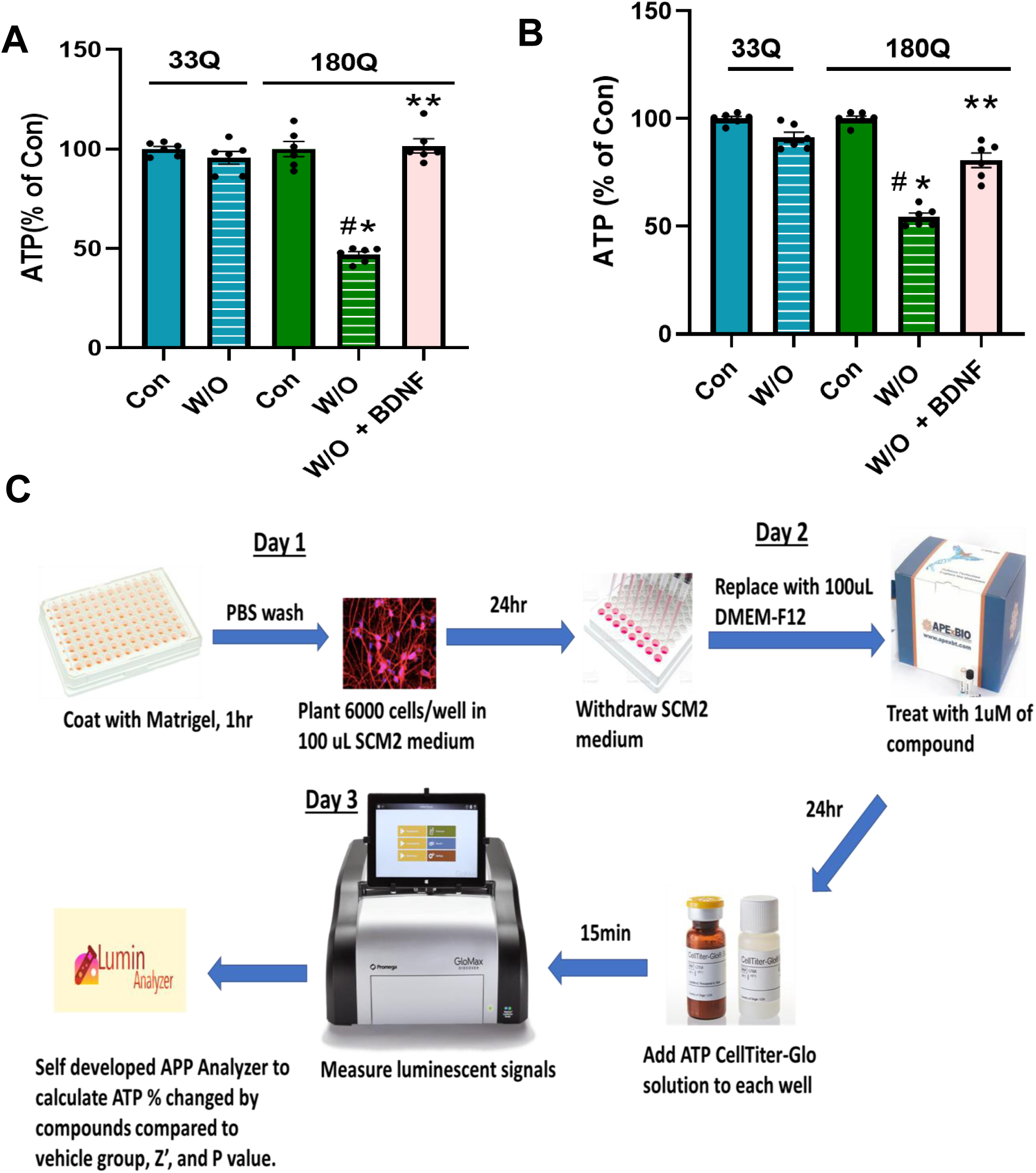
Drug screening platform utilizing HD striatal precursor neurons derived from human iPSCs. **A&B**) ISPNs with 33/18 and 180/18 CAG repeats were differentiated for 14 days (**A**) or maintained without differentiation (**B**), followed by stress induction via growth factor and nutrient withdrawal (W/O) for 24 hours. BDNF was added at the same time as cellular stress. ATP levels were quantified using the CellTiter-Glo cell viability assay. Results are expressed as MEAN ± SEM and normalized to the control (Con) group under normal growth conditions (n = 5–6). Statistical significance is analyzed by one-way ANOVA. *p < 0.05 vs. 33Q W/O, #p < 0.05 vs. 180Q Con, and **p < 0.05 vs. 180Q W/O. **C**) Schematic of the screening assay. ISPNs were seeded at a density of 6,000 cells per well in a 96-well plate format. Following 24 hours of growth factor and nutrient withdrawal and compound treatment, ATP levels were measured using luminescence-based detection with the GloMAX luminometer. Data analysis was performed using a custom-developed Lumin Analyzer application.

Additionally, we developed a custom application, Lumin Analyzer, to streamline assay result calculations. The Lumin Analyzer is a Python-based tool designed to support the analysis of high-throughput ATP assay in this study. It integrates raw output from luminometer with kinase library chemical databases, enabling automated statistical calculations and structured data visualization. Built using Python bindings for the Qt application framework, the tool supports real-time interaction with experimental data and accommodates flexible assay layouts. By replacing manual data handling with scripted workflows, the Lumin Analyzer reduces the potential for human error and improves consistency in analysis across experiments.

### Kinase inhibitor library screening identified 5 hits in HD striatal precursor neurons derived from human iPSCs

Kinases play critical roles in cellular signaling and are frequently dysregulated in various diseases, making the screening of kinase inhibitor libraries a widely used strategy in drug discovery and research ^24,25^. Additionally, our previous studies, along with findings from other researchers, suggest that HTT protein phosphorylation can modulate mHTT toxicity ^26–31^. We selected a kinase inhibitor library consisting of 764 inhibitors from ApexBio (Catalog No. L1024). The compounds are available as lyophilized powder or pre-dissolved in DMSO and are provided in a 96-well format sample storage tube with a screw cap and an optional 2D barcode. Since precursor ISPNs share a similar vulnerability to mHTT toxicity as mature neurons, we utilized precursors (with 180 CAG repeats) for initial screens.

We tested 764 compounds from the kinase inhibitor library over three rounds of screening. In the first round, all inhibitors were tested at a concentration of 1 µM. For additional quality control we have calculated Z’ factor for each experimental plate, and only plates with Z’ score >0.5 were included in the analysis ^10^. Additional screening criteria included a significant increase in ATP levels compared to vehicle control, and a minimum of 15% ATP increase over the vehicle control (Figure 2A). Hits from the first round were then subjected to dose-response testing in the second round. Finally, EC50 values were calculated for compounds demonstrating dose-dependent effects in the second round (Figure 2B). The overall results are summarized in Figure 2C, with the initial 23 hits identified in the first round listed in Supplementary Table 1. Among the 764 kinase inhibitors, most compounds (689) didn’t have effect on the ATP levels of the 180Q ISPNs. 52 compounds were toxic to the cells. From the initial 23 hits identified in the first round, we narrowed the selection down to 5 hits after completing all three rounds of screening. The final hits include the CK1 inhibitor D4476, ATR/CDK inhibitor NU6027, CDK1 and GSK3β inhibitor Kenpaullone, GSK3β inhibitor AZD1080, and PKC-α/β1 inhibitor GO6976. The detailed EC50 values for these 5 hits are presented in Figure 3A-E.

**Figure 2.**
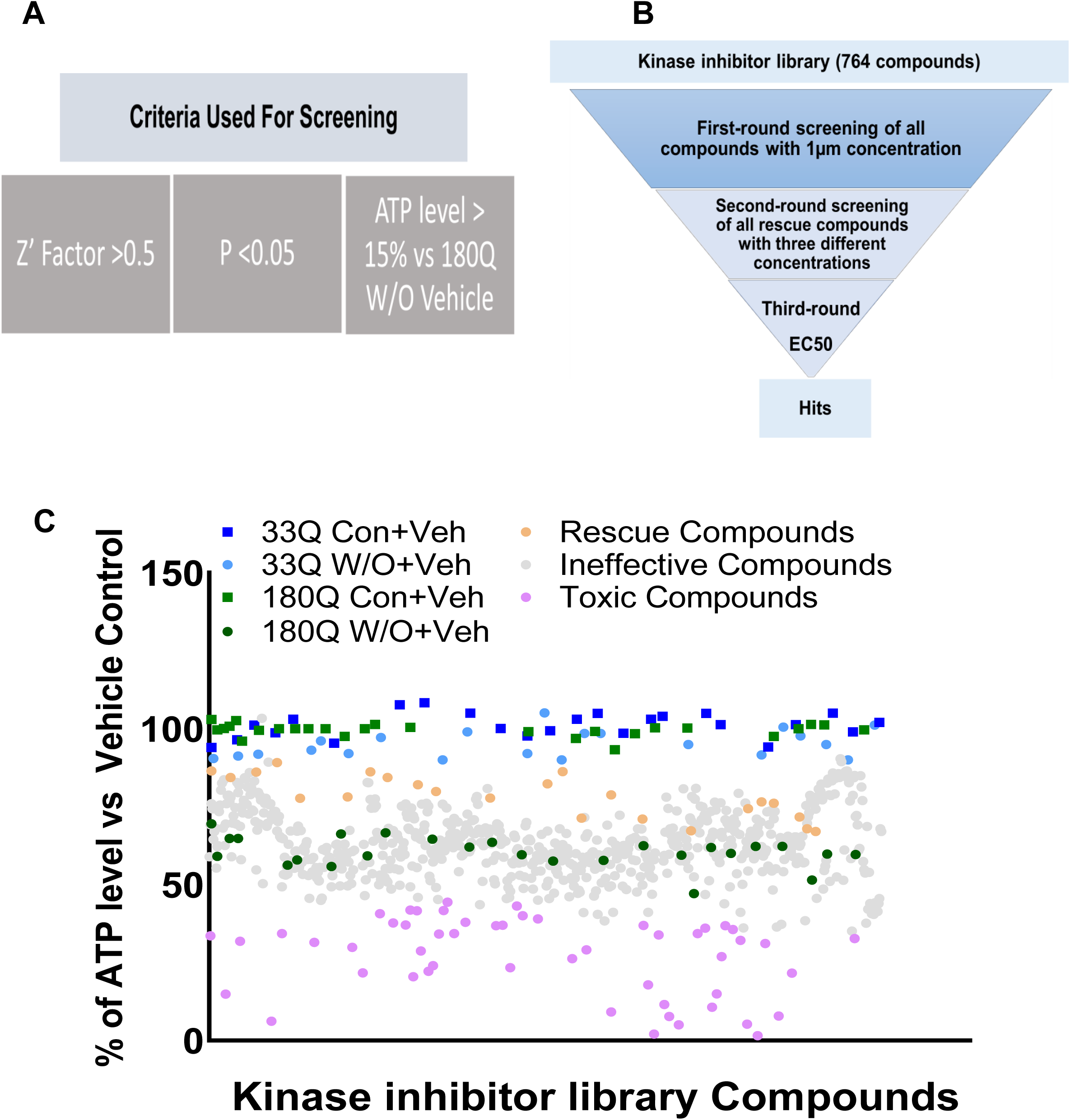
Kinase inhibitor library screening in HD striatal precursor neurons derived from human iPSCs. **A).** Screening Criteria: Plates required a Z’ factor > 0.5, a significant ATP increase compared to the vehicle control, and at least a 15% ATP increase over the vehicle control. **B).** A library of 764 kinase inhibitors was screened in three stages. In the first round, all compounds were screened at 1 µM. Hits advanced to the second round for dose-response testing. EC50 values were determined in the final round for compounds showing dose-dependent effects. **C).** Overview of screening results from first round.

**Figure 3.**
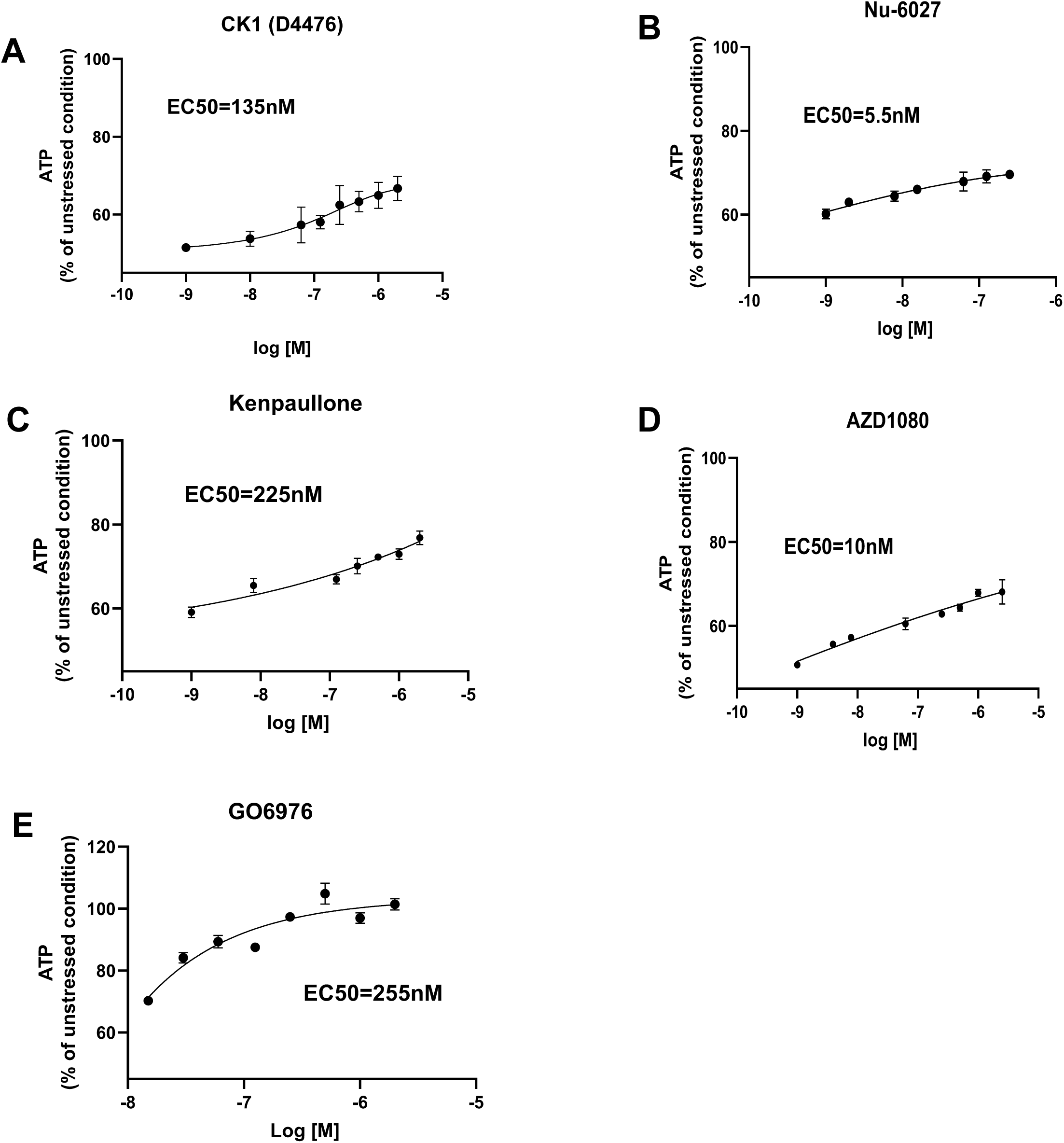
EC50 values for 5 top hits from a kinase inhibitor library screen in HD striatal precursor neurons derived from human iPSCs. Detailed EC50 values for the hits are provided (**A-E**). The top hits were CK1 inhibitor D4476, ATR/CDK inhibitor NU6027, CDK1/GSK3β inhibitor Kenpaullone, GSK3β inhibitor AZD1080, and PKC-α/β1 inhibitor GO6976.

### PKC-a& β1 inhibitor GO6976 protects HD human striatal precursor neurons against stress-induced toxicity

Among the 5 hits, one compound, GO6976 (Figure 4A), provided the greatest effect on ATP levels and dose-dependently protected HD striatal precursor neurons (Figure 4B). GO6976 was initially characterized as a potent and selective inhibitor of conventional PKC isoforms (α and β1)^32^. To further investigate the specificity of neuroprotection by PKC inhibitors, we tested the pan-PKC inhibitor GO6983, the PKC-α inhibitor Tetrandrine ^33^, and a PKC-β inhibitor LY333531 ^34^. Notably, only GO6976 provided protection (Figure 4C), suggesting that simultaneous inhibition of both PKC-α and PKC-β1 is required for the observed protective effect. We confirmed the protective effect of GO6976 in HD mature striatal neurons differentiated from ISPNs (Supplementary Figure 2). GO6976 increases ATP levels in ISPNs with 33 CAG repeats under cellular stress, surpassing even unstressed conditions (Supplementary Figure 3). This suggests that GO6976 enhances the overall health of the cells. To rule out any artificial interaction between GO6976 and the assay reagent, we validated the results using the CellTiter-Glo luminescent assay reagent alone, without cells, as shown in Supplementary Figure 4.

**Figure 4.**
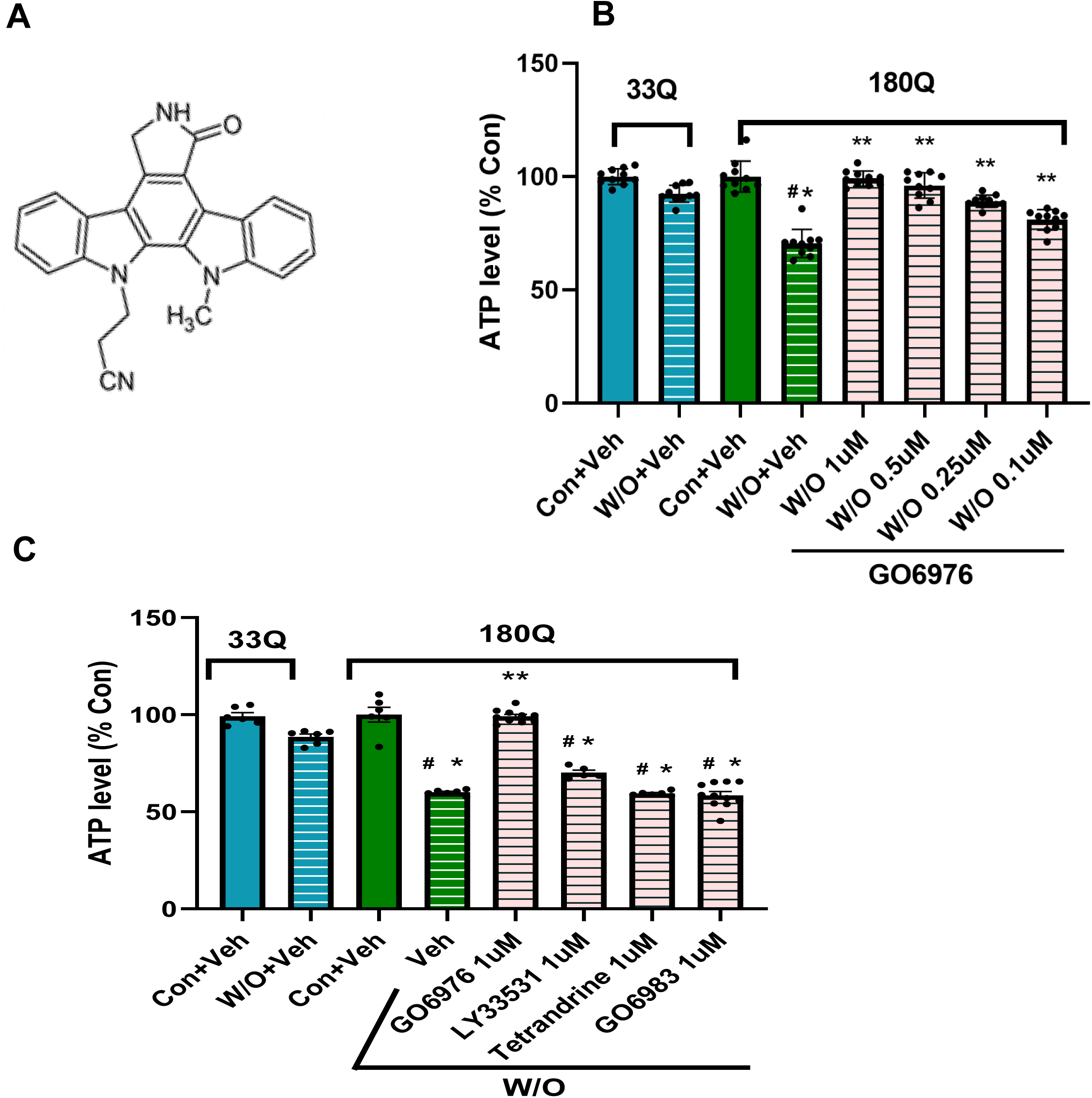
PKC-a&β1 inhibitor GO6976 protects HD striatal precursor neurons against stress-induced toxicity in a dose-dependent manner. **A).** GO6976 structure formulas. **B).** ISPNs with 33/18 and 180/18 CAG repeats were stressed by growth factor and nutrients withdrawal (W/O) for 24 hr. 180Q ISPNs were treated with GO6976 at 0.1,0.25,0.5, 1 uM, or Veh (Vehicle control). **C).** 180Q ISPNs were treated with 1uM GO6976, LY333531, Tetrandrine, GO6983 or Veh upon stress. ATP levels were measured using CellTiter-Glo cell viability assay. The results from B&C are presented as MEAN ± SEM, normalized as % of the value of control (Con) group under normal growth conditions (n=5-10). One-way ANOVA, * p<0.05 vs 33Q W/O+Veh; # p<0.05 vs 180Q Con+Veh; ** p<0.05 vs 180Q W/O+Veh.

### Genetic modulation of PKC-a& β1 expression regulates mutant Huntingtin toxicity

GO6976, developed by GÖDECKE AG (later acquired by Pfizer) in the early 1990s, is a potent and selective inhibitor of conventional Ca^2+^-dependent PKC isoforms (α and β1) ^32^. It was later widely used in academic research to study PKC-mediated signaling, particularly in cancer models such as acute leukemia and bladder cancer ^35,36^. Subsequently, GO6976 was found to inhibit other kinases as well ^36–39^.

To investigate the specificity of PKC-α and PKC-β1 in modulating mHTT toxicity, we overexpressed PKC-α, PKC-β1, or both, in 180Q ISPNs via electroporation (Figure 5A). Overexpression of both PKC-α and PKC-β1 together resulted in significant toxicity in 180Q ISPNs, whereas overexpression of either gene alone did not (Figure 5B). In contrast, knockdown of PKC-α and PKC-β1 conferred partial protection (Figures 5C and 5D). These results indicate that the combined activity of PKC-α and PKC-β1 contributes to mHTT-induced toxicity and that their modulation may offer a therapeutic strategy.

**Figure 5.**
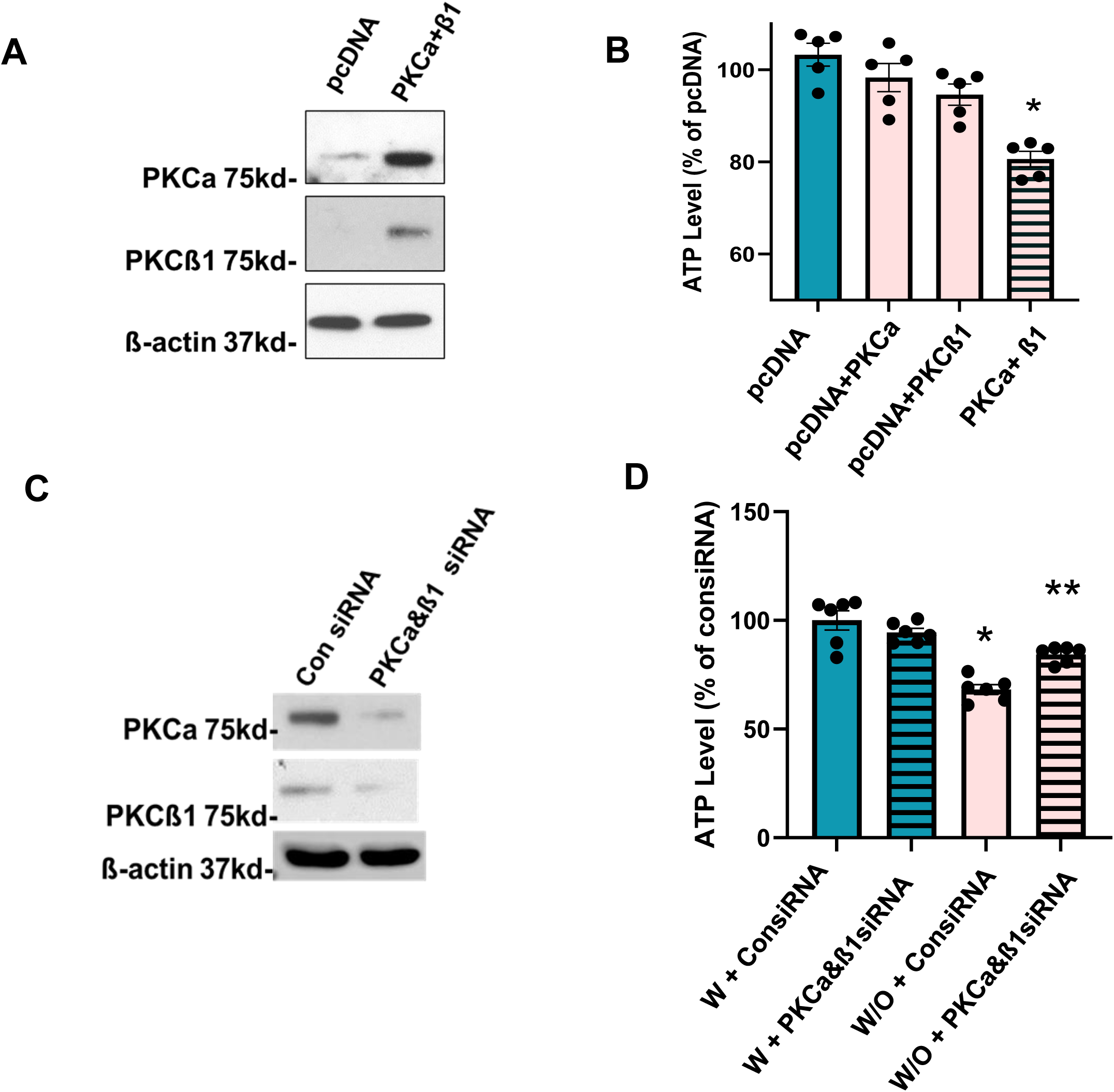
Manipulation of PKC-a&β1 expression alters mutant Huntingtin toxicity. **A & B**) Co-expression of PKC-α and PKC-β1 induces cellular toxicity in HD ISPNs. 180Q ISPNs were electroporated with pcDNA (control), PKC-a, PKC-β1, or a combination of PKC-α and PKC-β1. Cell lysates were collected 24 hrs post-electroporation and analyzed by immunoblotting for PKC-α and PKC-β1 expression. **A**). ATP levels were quantified using the CellTiter-Glo viability assay 24 hours post-electroporation. **B**). Data are presented as mean ± SEM, normalized to the pcDNA control (n=5). One-way ANOVA, *p < 0.05 compared to pcDNA. **C & D**) Knockdown of PKC-α and PKC-β1 alleviates mHTT-induced cellular toxicity in HD ISPNs. 180Q ISPNs were electroporated with either control siRNA or a combination of PKC-α and PKC-β1 siRNA for 48 hrs, then stressed by growth factor and nutrients withdrawal (W/O) for 24 hrs. Cell lysates were collected and analyzed by immunoblotting to confirm the knockdown of PKC-α and PKC-β1 (**C**). ATP levels were measured using the CellTiter-Glo viability assay 72 hours post-electroporation. Data are presented as mean ± SEM, normalized to the normal growth condition with control siRNA (W+ConsiRNA) (n=6). One-way ANOVA, *p < 0.05 compared to W+consiRNA, **p<0.05 compared to W/O+consiRNA (D).

### Increased phosphorylation of conventional PKC-a and PKC-β1 in HD models

PKC is a diverse family of enzymes that transduce a large variety of cellular signals. In the brain, PKC is critical for regulating synaptic plasticity, learning and memory. PKC plays a key role in controlling the balance between cell survival and cell death ^40^. Increase of PKC activity is associated with neurodegeneration ^41^. There is multiple evidence that phosphorylation at a number of sites controls the activity of PKC α/ β1 (reviewed in ^42^). To investigate whether PKC-α and PKC-β1 are involved in HD pathogenesis, we assessed their phosphorylation state in HD cell and animal models. We first evaluated phosphorylated PKC-α and PKC-β1 in our ISPNs with either 33 or 180 CAG repeats. Several serine and threonine phosphorylation sites were tested, and we found a significant increase in PKC-α and β1 phosphorylation in 180Q HD ISPNs compared to 33Q controls, specifically at S657 on PKC-α and T642/644 on PKC-β1, sites known to regulate the activity of PKC α/ β1^42^ (Figures 6A-C). These findings were further confirmed in postmortem human HD brains (Figures 6D-F) and 250Q HD mouse striatum tissues (Figures 6G-I). Notably, total PKC-β1 expression showed variability among postmortem HD human brains, with levels generally decreased compared to controls. However, when normalized to total PKC-β1, phosphorylation levels were consistently elevated in all HD cases.

**Figure 6.**
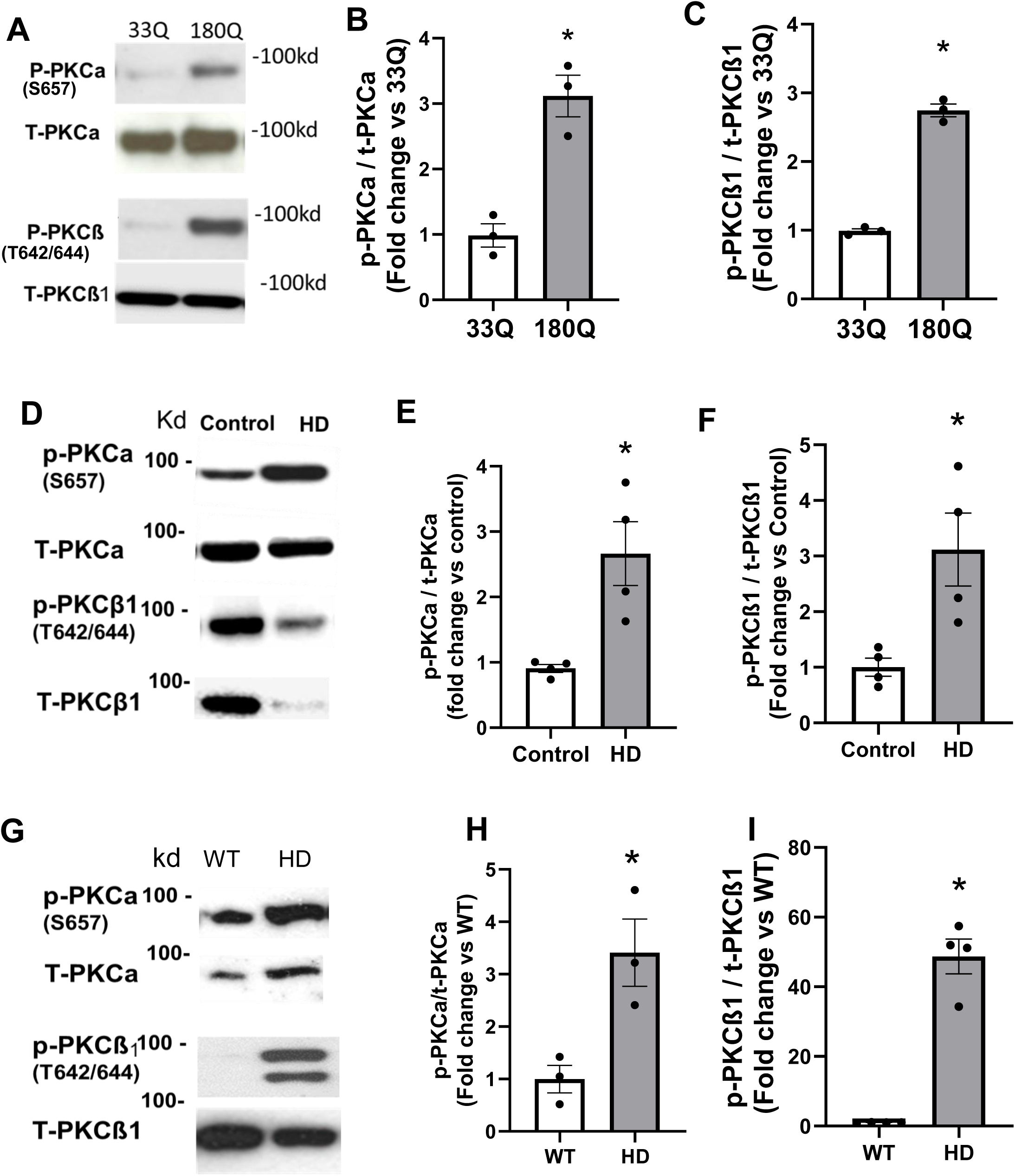
The phosphorylation of conventional PKC-a and PKC-β1 increased in HD ISPNs, HD mouse and HD human post-mortem brains. **A-C**). The phosphorylation of PKC-a at S657 and PKC-β1 at T642/644 increased in HD ISPNs after cellular stress for half hour. 33Q or 180Q ISPNs were stressed by withdrawal of growth factors and nutrition for 1/2h. Western blotting was performed to detect phosphorylated PKC-a (p-PKCa), total PKC-a (t-PKCa), p-PKCβ1, or t-PKCβ1 in ISPNs. **D-F**). The phosphorylation of PKC-a and PKC-β1 increased in HD postmortem human brain motor cortex BA4 tissues. Four cases of BA4 tissues from HD patients (Vonsattel grade II-IV) or age matched controls were lysed to perform western blot to detect phosphorylated and total PKC-a or PKC-β1. **G-I**). The phosphorylation of PKC-a at S657 and PKC-β1 at T642/644 increased in HD ZQ250 mouse striatal tissues at age of 6 months. WT or HD striatum tissues from our lab brain bank were lysed and subjected to western blot analysis to detect both phosphorylated and total protein levels of PKC-a or PKC-β1. All blots were quantified by NIH Image J software. *P < 0.05 by Student’s t-test. The results are presented as MEAN ± SEM, normalized as % of the value of control (n=3-4). * p<0.05 vs control.

PKC isoforms are categorized into conventional, novel, and atypical groups. PKC-α and β1 belong to the conventional PKCs, which require the second messengers diacylglycerol (DG) and Ca²⁺ for activation ^43^.To determine whether the activity changes in HD samples were specific to conventional PKC-α and β1, we assessed non-conventional PKCs. Both the activity and total levels of PKC-δ and PKC-θ were decreased in HD ISPNs (Figures 7A-E). These results highlight a specific increase in PKC-α and PKC-β1 activity in HD, which may explain why pan-PKC inhibitors fail to protect HD neurons.

**Figure 7.**
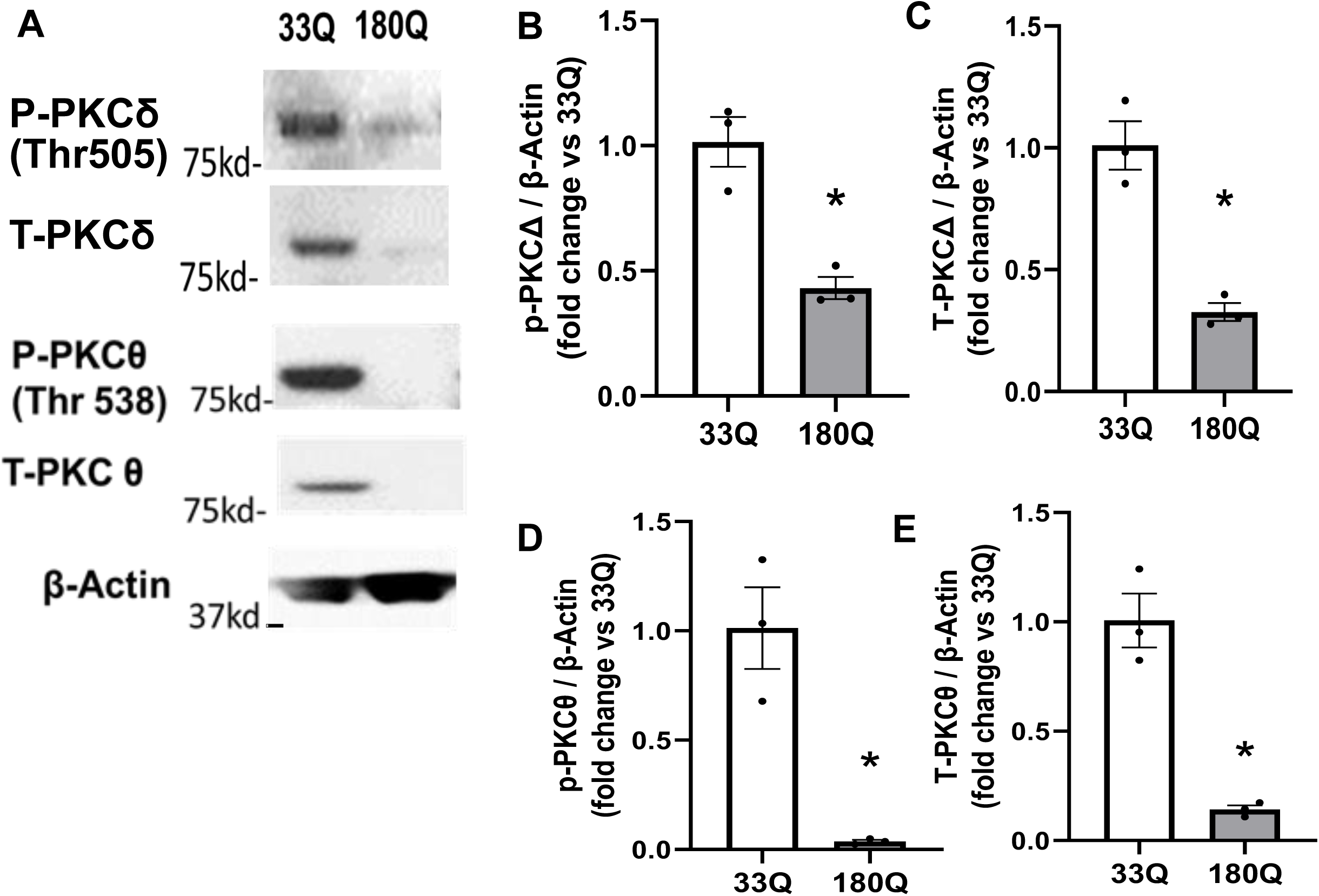
The phosphorylation and levels of non-conventional PKC-δ and PKC-θ were decreased in HD ISPNs. 33Q or 180Q ISPNs were stressed by withdrawal of growth factors and nutrition for ½ h. Western blotting was performed to detect phosphorylated PKC-δ (p-PKC δ), total PKC-δ (t-PKCδ), p-PKCθ, or t-PKCθ (**A**). Blots were quantified by NIH Image J software. *P < 0.05 vs 33Q by t-test. The results are presented as MEAN ± SEM, normalized as % of the value of 33Q (n=3) (**B-E**).

### PKC-a and PKC-β1 Interact and colocalize with wild-type and mutant HTT

To further investigate how inhibiting PKC-α and PKC-β1 protects HD neurons, we overexpressed full-length wild-type or mutant HTT along with PKC-α or PKC-β1 in 293T cells and performed pull-down assays to assess interactions between PKC-α/β1 and HTT (Figures 8A-B). We found that both PKC-α and PKC-β1 associate with wild-type and mutant HTT. Notably, endogenous interaction between HTT and PKC-β1 was detected in 293T cells transfected with PKC-β1 alone. The physiological co-association of wild-type and mutant HTT with PKC-α or PKC-β1 was further validated in human postmortem control and HD brain tissues (Figure 8C).

**Figure 8.**
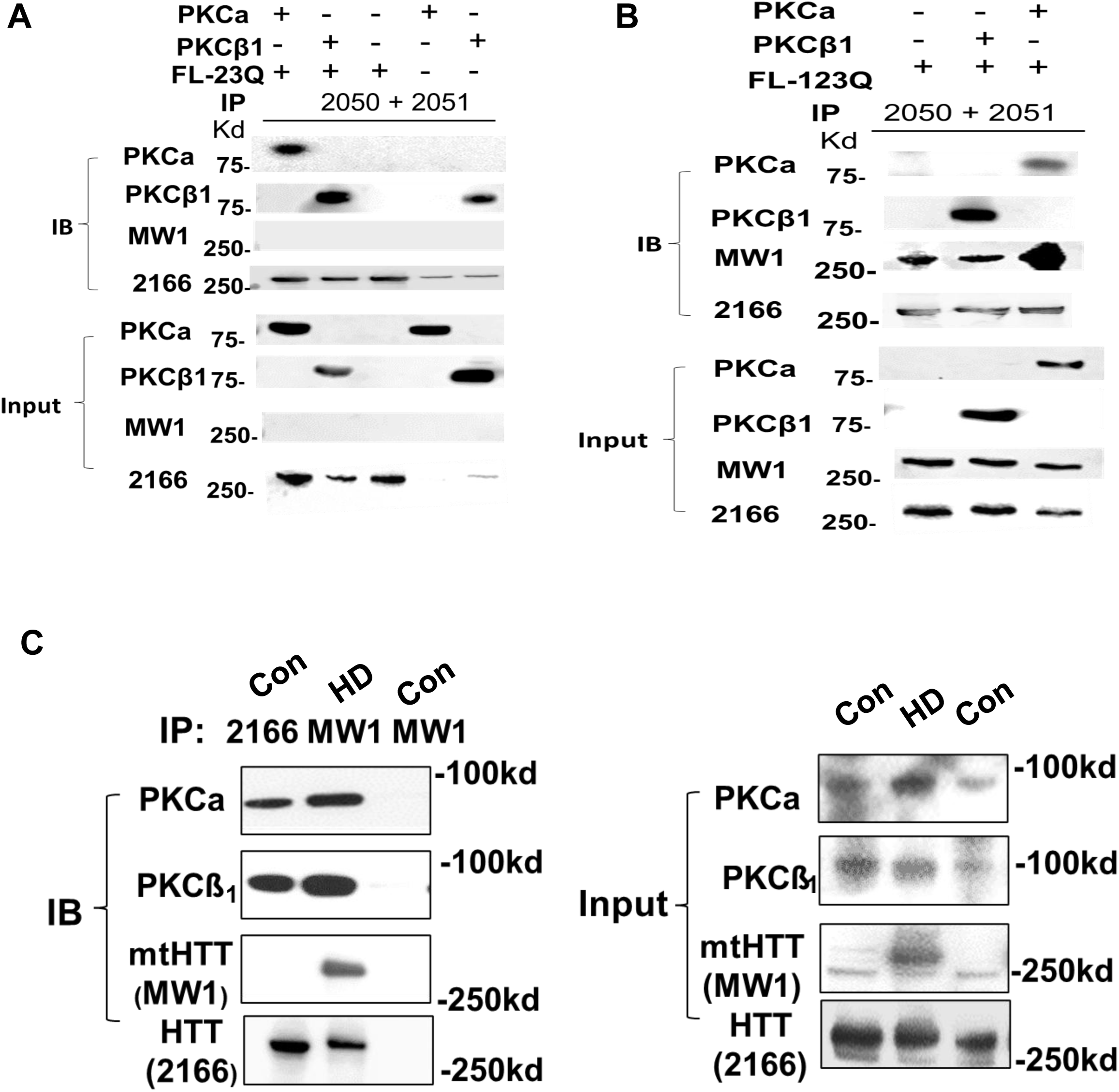
PKC-a and PKC-β1 interact with wildtype and mutant HTT. **A&B**). HEK293-FT cells were co-transfected with full length HTT with 23Q (wHTT) or 123Q (mHTT) and PKC-a or PKC-β1 respectively for 24 h. Cell lysates were subjected to co-IP assays using HTT antibody (MCA2050 and 2051 to pull down both cytosolic and nuclear HTT), followed by anti-PKC-a, anti-PKC-β1, anti-HTT (2166 antibody), or anti-polyglutamine (mHTT, MW1) immunoblotting. **C**). Cortical brain homogenates from age-matched controls and HD patients were subjected to co-IP assays with anti-HTT or anti-polyglutamine antibodies followed by anti-PKC-a or PKC-β1 immunoblotting.

Immunostaining and confocal imaging showed that PKC-α predominantly colocalizes with HTT in the cytoplasm, while PKC-β1 colocalizes with HTT in the nucleus (Figure 9A). However, proximity ligation assays (PLA) revealed that the interactions between HTT and PKC-α/β1 occur in both cytoplasmic and nuclear compartments (Figure 9B).

**Figure 9.**
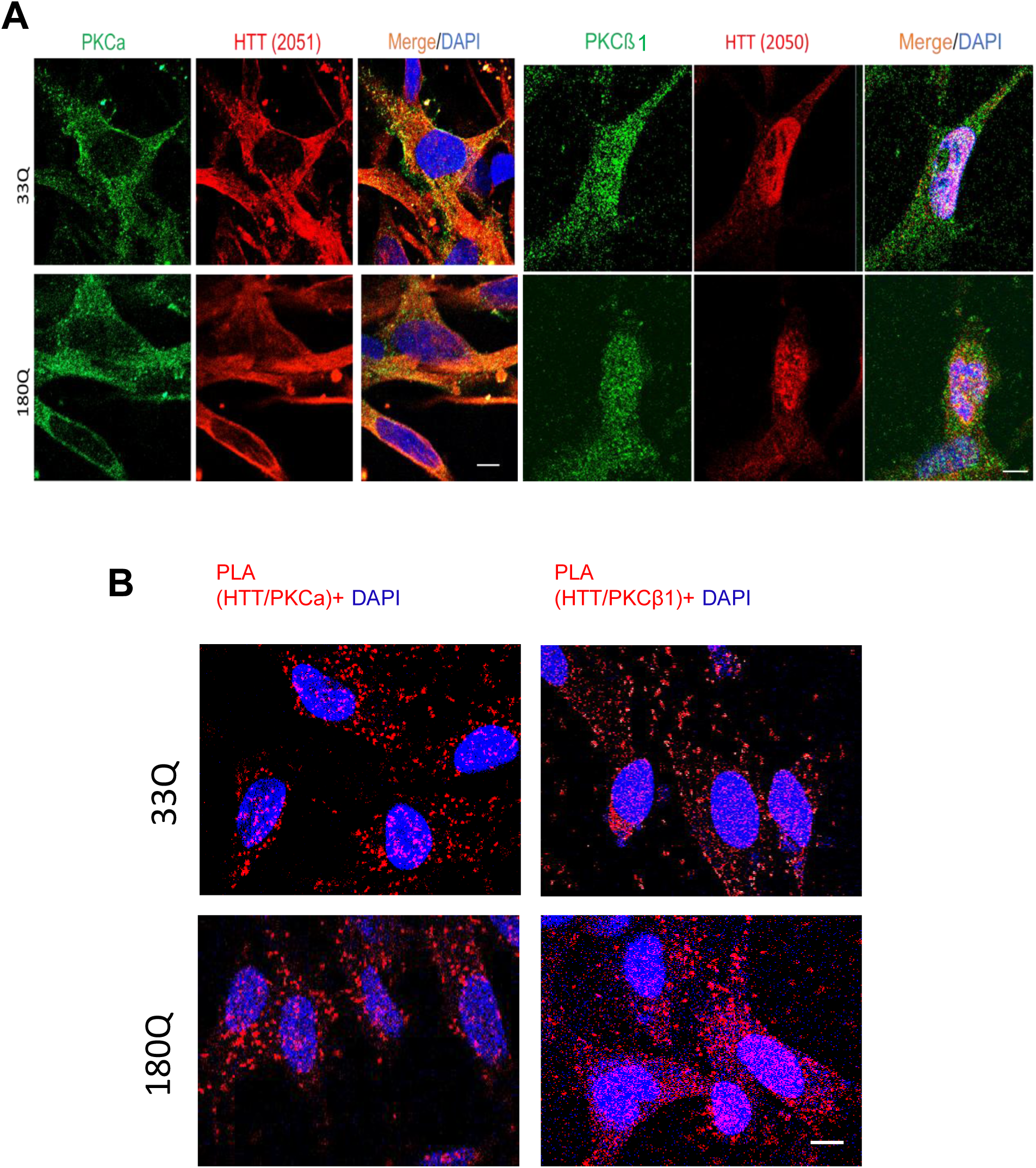
PKC-α and PKC-β1 colocalize with wild-type and mutant HTT in both the cytosolic and nuclear compartments of ISPNs. A). Human striatal precursor neurons from normal human subjects with 33Q or HD patients with 180Q were subjected to co-immunostaining using anti-HTT (MCA2050/2051, red) and anti-PKC-a/PKC-β1 (green) antibodies. Scale bar 5 uM. B). Human striatal precursor neurons from normal human subjects with 33Q or HD patients with 180Q were subjected to in situ interaction assay using Duolink in situ red kit. PLA signals are shown in red and the nuclei in blue. Scale bar 5 uM.

## Discussion

In this study, we developed a screening platform based on human striatal neurons to evaluate potential therapeutics targeting mutant huntingtin toxicity. Our findings demonstrate that ISPNs derived from HD iPSCs provide a robust cellular model for drug screening and target validation in Huntington’s disease. Furthermore, we identified a small-molecule PKC-α/β1 inhibitor as a potential candidate for HD treatment and validated that inhibiting PKC-α and β1 offers protective effects in HD.

Human iPSCs offer significant advantages over traditional cell models, particularly for studying genetic disorders ^44^. By reprogramming somatic cells from patients, iPSCs enable the generation of scalable and disease-relevant human cells. These patient-specific cellular models have become instrumental in drug screening efforts, providing a powerful platform for identifying and validating novel therapeutics tailored to human disease ^45^. The traditional iPSC neuronal differentiation process posed significant challenges for generating large quantities of cells required for biochemical or screening assays. Our ISPNs overcame this limitation, providing a valuable platform for drug screening in human striatal neurons. For immortalization, we utilized co-expression of the enzymatic component of telomerase (hTERT) to maintain genomic stability and conditional expression of c-Myc fused to the tamoxifen-sensitive hormone-binding domain of the estrogen receptor (ER). The regulation of c-Myc-ER expression via tamoxifen withdrawal enables gene silencing during compound library screening, neuronal differentiation, and experimental assays, thereby preventing potential disruptions caused by constitutive expression of immortalizing proto-oncogenes ^23^.

Our initial screening of 764 compounds from the kinase inhibitor library identified 23 hits. Subsequent second- and third-round rescreens prioritized 5 compounds, yielding a final hit rate of 0.65%, which falls within the acceptable range and indicates that both the assay and compound library performed as expected. We focused our follow-up studies on PKC inhibitors, as GO6976 showed the strongest protective effect among the identified hits. Among the four other identified hits, CK1, CDK, and GSK emerged as potential therapeutic targets warranting further validation. CDK and GSK have been extensively studied in the context of HD, and their roles in disease progression and therapeutic modulation are well understood ^30,46–48^. In contrast, the potential involvement of CK1 in HD pathology remains largely unexplored. CK1 is a serine/threonine kinase family with multiple isoforms, each playing distinct roles in cellular processes such as DNA repair, circadian rhythm regulation, and Wnt signaling ^49^. Given the critical role of these pathways in neuronal survival and function, investigating CK1 isoforms in HD could provide new insights into disease mechanisms and therapeutic opportunities. Further studies are necessary to elucidate the specific contributions of CK1 to HD pathophysiology, including its potential as a modulator of mutant huntingtin toxicity. This line of research could open avenues for the development of isoform-specific inhibitors or modulators, offering a novel strategy for HD treatment.

PKC is classified into three categories: conventional, novel, and atypical. Dysregulation of PKC activity has been implicated in numerous diseases, including cancer, cardiovascular disorders, and particularly neurodegenerative conditions. Gain-of-function mutations in PKCα have been linked to Alzheimer’s disease (AD), and mutations in PKC-α increase activity in Alzheimer’s disease cases. Thus, elevation of PKC-a signaling is an early event in Alzheimer’s disease pathogenesis ^41,50^. These findings underscore the involvement of PKC in the pathogenesis of neurodegenerative diseases. In HD, PKC signaling has also been implicated. Previous studies reported decreased PKC-βII immunoreactivity and mRNA levels in the caudate-putamen of HD patients and HD mouse striatum tissues ^51,52^. Additionally, PKC-δ protein levels were reduced in the striatum and cortex of R6/1, R6/2, and Hdh(Q111/Q111) mice, as well as in the putamen of HD patients ^53,54^. PKC-δ was specifically associated with intranuclear aggregates in R6/2 mice and in Neuro2a cells expressing N-terminal truncated huntingtin with 150Q ^53^. These findings highlight the involvement of PKC in HD pathology. However, despite these insights, follow-up research on PKC as a therapeutic target—particularly its kinase activity, has been limited. We observed a significant decrease in the activity and expression of non-conventional PKCs, including PKC-δ and PKC-θ in HD ISPNs. This finding is consistent with Rue’s results, which demonstrated similar reductions in PKC-δ expression across various HD models ^54^. In contrast, Zemskov’s studies in HD transgenic models reported nuclear accumulation of non-conventional PKC-δ^53^, suggesting model-specific differences in PKC regulation and localization. These discrepancies highlight the need for a more detailed investigation into the subcellular localization and activity of PKC-δ and PKC-θ to uncover their roles in HD pathology.

In this study, we identified GO6976, a PKC-α and PKC-β1 inhibitor, as the most protective compound from a library of over 750 kinase inhibitors. Although a limitation of our study is the lack of additional inhibitors targeting the same PKC-α/β1 subclass for independent validation, we confirmed that PKC-α and PKC-β1 activities are elevated in samples expressing endogenous full-length mHTT. Furthermore, we demonstrated that modulating PKC-α and PKC-β1 activity can regulate mHTT-induced toxicity. These findings suggest that PKC-α and PKC-β1 are promising therapeutic targets for further in vivo evaluation in HD mouse models.

Since PKC-α and PKC-β1 interact with huntingtin, the increased phosphorylation of these kinases may be directly driven by mHTT. Further mechanistic studies are needed to determine this point of view. Notably, intrathecal administration of GO6976 to 5xFAD mice restored alterations in the Y-maze test and ameliorated DNA double-strand breaks^55^. This suggests that modulating DNA damage and repair pathways could represent an additional mechanism of action of GO6976 worth further investigation, especially given their prominence in current HD research. Recent studies have demonstrated that mismatch DNA repair plays a critical role as a modifier of HD onset and a driver of CAG repeats somatic instability ^56,57^. Furthermore, our concurrent study highlights the pivotal role of the HTT protein in regulating interconnected DNA repair and remodeling pathways ^58^. Most importantly, further efficacy and mechanistic studies of GO6976 in HD mice will advance its development as a potential therapeutic for HD.

In conclusion, our study highlights the potential of a small-molecule PKC-α and PKC-β1 kinase inhibitor as a therapeutic strategy to mitigate mutant HTT toxicity in HD MSNs. By specifically targeting these kinases, our findings suggest a novel approach to reducing cellular dysfunction associated with HD pathology. Our work underscores the value of human iPSC-derived neuronal models as a robust and physiologically relevant platform for drug discovery. These models recapitulate key aspects of the disease, enabling the identification and validation of potential therapeutic candidates. Further preclinical investigation of PKC-α and PKC-β1 inhibitors may pave the way for novel treatments for HD.

## Supporting information

All supplemental files

## RESOURCE AVAILABILITY

### Lead contact

Further information and requests for resources and reagents should be directed to and will be fulfilled by the lead contact, Dr. Mali Jiang

### Materials availability

This study did not generate new unique reagents.

### Data and code availability

- Kinase inhibitor screening data and other data reported in this paper will be shared by the lead contact upon request.
- This paper does not report original code for Lumin Analyzer.
- Any additional information required to reanalyze the data reported in this paper is available from the lead contact upon request.

## ACKNOWLEDGMENTS

We thank Hoku West-Foyle from the JHMI Microscope Facility for providing technical support with confocal imaging analysis.

This work was supported by NINDS grant NS129563 (M.J.); NS104320, NS086452 (C.R.); NS135139, NS124084 (W.D.); P30AG0066507 (J.T.).

PKC-a (Addgene plasmid #21232) and PKC-β1 (Addgene plasmid #16378) are gifts from Dr. Bernard Weinstein (Soh and Weinstein 2003).

## AUTHOR CONTRIBUTIONS

M.J. and C.R. designed research. T.S., R.M., M.R., R.W., R.S., L.G., J.J., K.B., Y.L., A.C., K.B., Y.U., A.Y., C.H., J.T. performed experiments, analyzed data. Y.X. designed and coded the Lumin Analyzer App. J.T. provided human tissues, W.D. provided HD mouse tissues, T.R. and W.S. analyzed data. M.J., L.M., and T.R. designed the graphic abstract. M.J., C.R., W.S., and T.R. wrote the paper.

## STAR★ Methods

### Key resources table

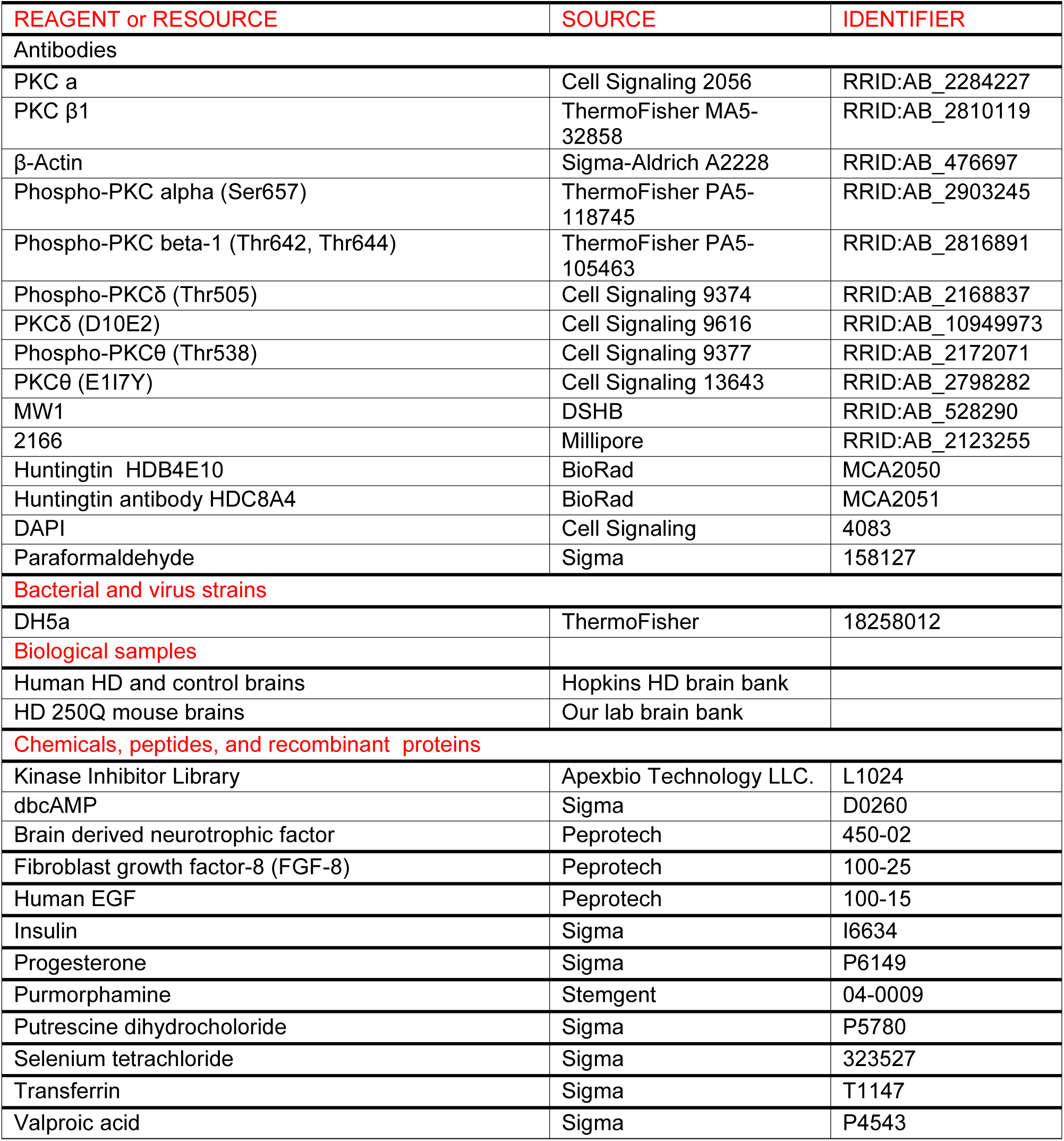
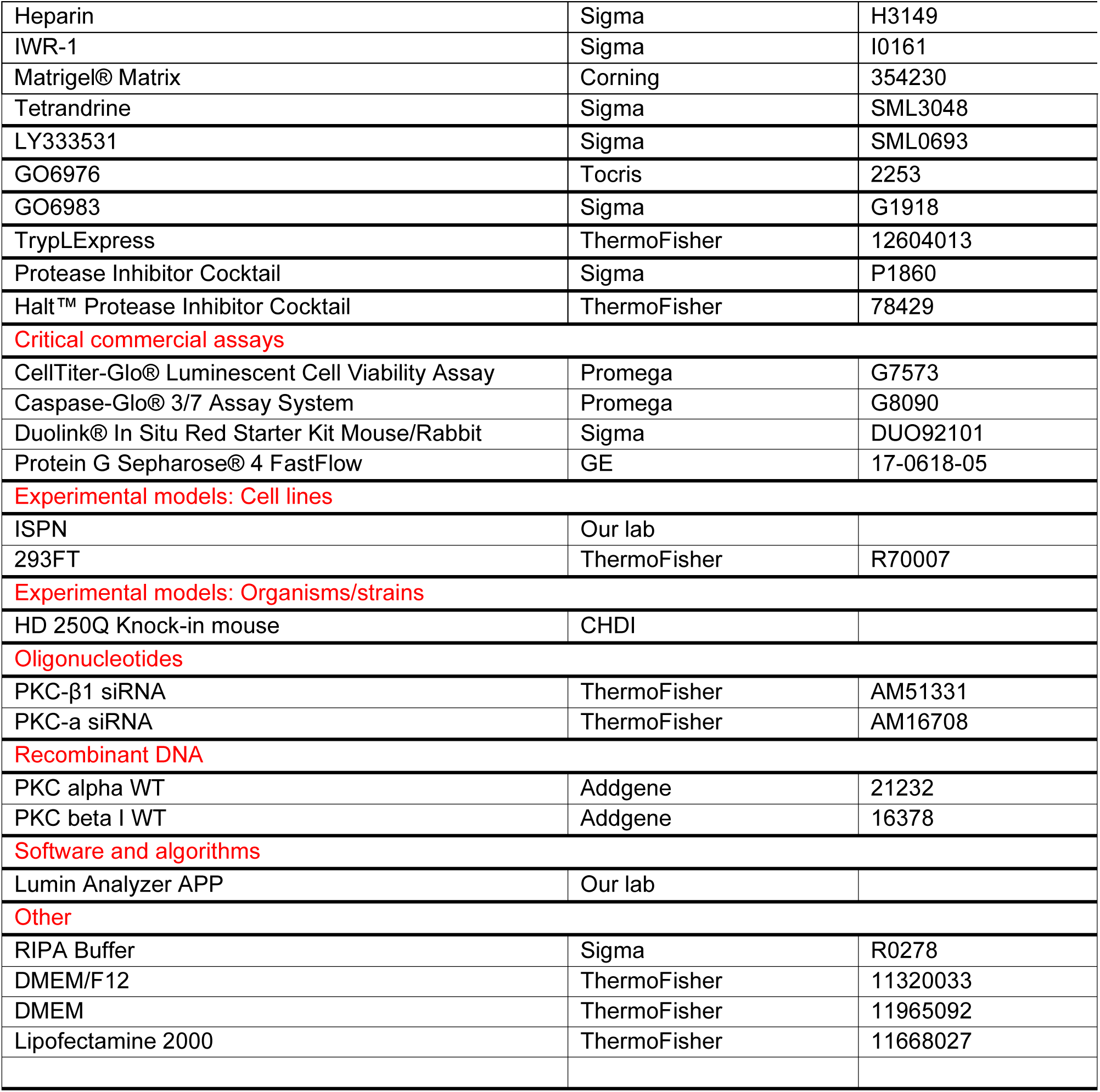

### Method details

#### Cell culture

Immortalized Striatal Precursor Neurons (ISPNs) were generated in our laboratory from patient-derived iPS cells carrying either normal (33) or expanded (180) CAG repeats, using co-expression of the enzymatic component of telomerase (hTERT) and drug-regulated c-Myc, as previously described ^23^. Notably, the 180-CAG HD ISPN line has undergone progressive CAG repeat expansion, with the repeat size ranging from 240 to 260 at the time of these studies. For consistency, we refer to this line as 180-CAG throughout this paper.

The ISPNs were maintained as stable, adherent neuronal precursors on Matrigel-coated plates in SCM2 medium at 37°C with 5% CO₂. Differentiation to a phenotype resembling medium spiny neurons was performed as previously described ^23^.

Human embryonic kidney (HEK) 293FT cells, obtained from Invitrogen (Thermo Fisher), were cultured in DMEM (containing 4.5 g/L D-glucose; ThermoFisher) supplemented with 10% FBS, 100 µg/ml Geneticin, 100 U/ml penicillin, and 100 U/ml streptomycin.

#### Cell viability assays

ISPNs were plated at a density of 6,000 cells per well in 96-well plates. Stress was induced by growth factor and nutrient withdrawal for 24 hours, followed by treatment with test compounds. ATP levels were measured using the CellTiter-Glo® Cell Viability Assay and quantified with a GloMAX® luminometer ^23^.

#### The Development of the Lumin Analyzer

The Lumin Analyzer is a Python-based tool designed to automate data analysis, streamline workflows, and generate reports efficiently. It simplifies complex and repetitive tasks, such as integrating instrument-generated Excel reports with compound chemical databases, calculating statistics, and visualizing results.

The Workflow is as follow: 1). Upload raw experimental data and chemical database. 2). Define group structures for analysis (e.g., treatment vs. control). 3). Process data to compute statistics and generate subgroup results. 4). Visualize results with dynamic charts and tables. 5). Generate a finalized report linking experimental and chemical data.

The Lumin Analyzer significantly accelerates data processing, reduces errors, and provides a user-friendly platform for high-throughput analysis and reporting.

The code for the Lumin Analyzer will be made available to the public upon request.

#### Celetrix electroporation

ISPNs are trypsinized in TrypLExpress, counted, and resuspended in the proprietary Celetrix electroporation buffer. The 1×10^6^ cell suspension is mixed with 1ug pcDNA, PKC-a, PKC-β1, PKC-a+β1, or control siRNA, or 100nM PKC-a/β1 siRNA and transferred to the Celetrix 120ul tubes. Electroporation is performed using 640 V and 30 ms. Following electroporation, cells are immediately transferred to a pre-warmed culture medium and incubated under standard conditions for recovery and downstream analysis.

#### Protein extraction and western blot analysis

Cells and brain tissues were homogenized and lysed in 9 volumes of RIPA buffer (Sigma) containing 50 mM Tris–HCl (pH 8.0), 150 mM sodium chloride, 1.0% Igepal CA-630 (NP-40), 0.5% sodium deoxycholate, 0.1% sodium dodecyl sulfate (SDS), and freshly prepared protease inhibitor (1:100, Sigma) and/or phosphatase inhibitor (1:100, Sigma). The lysates were centrifuged at 14,000 × g for 15 minutes at 4°C, and the supernatant fractions were collected. Protein concentrations were determined using the Micro BCA™ Protein Assay Kit (Pierce Protein Research Products).

Soluble proteins were separated by SDS–PAGE and transferred onto a nitrocellulose membrane. The membrane was blocked with BSA or 5% nonfat milk and incubated overnight at 4°C with primary antibodies against PKC-α, PKC-β1, p-PKC-α, p-PKC-β1, PKC-δ, PKC-θ, p-PKC-δ, or p-PKC-θ. Subsequently, the membrane was incubated for 1 hour with an HRP-conjugated secondary antibody (1:3000; GE Healthcare). Immunoreactive proteins were visualized using a chemiluminescence detection kit following the manufacturer’s protocol (ECL Kit; Amersham Corp. or Supersignal West Pico PLUS Chemiluminescent Substrate, ThermoFisher).

#### Transfection

Lipofectamine 2000 (Invitrogen) was used to co-express full length wHTT or mHTT with PKC-a or PKC-β1 in human HEK293-FT cells. 10 μg plasmid DNA and 30 μl lipofectamine/10 cm dish were mixed. Cells were collected for western blotting, or immunoprecipitation (IP) at 24 h after transfection.

#### Immunoprecipitation

HEK293-FT cells or human brain tissues were lysed in RIPA buffer (Cell Signaling) and pre-cleaned by incubation with Protein G Sepharose® 4 Fast Flow (GE) for 1 hour. HTT was co-immunoprecipitated using MCA2050/2051 antibodies for 12 hours, followed by incubation with Protein G Sepharose® 4 Fast Flow for 2 hours. The beads were then washed extensively with RIPA buffer and PBS before elution. Solubilized proteins were separated by SDS–PAGE and analyzed by western blot using antibodies against PKCα, PKCβ1, MW1, and MAB2166.

#### Immunostaining

ISPNs were fixed by 4% paraformaldehyde for 10 mins and co-immunostained with HTT (2050/2051) and PKC-a or PKC-β1. Images were taken using Zeiss Confocal Florescence microscope.

#### Proximity Ligation Assay (PLA)

Culture ISPNs on glass coverslips and fix them with 4% paraformaldehyde. After permeabilizing with 0.1% Triton X-100, block non-specific binding with blocking buffer. Incubate the cells with anti-mouse HTT and anti-rabbit PKCα or PKCβ1 primary antibodies. Apply PLA probes (secondary antibodies conjugated to unique DNA oligonucleotides) specific to mouse and rabbit primary antibodies. If HTT and PKCα/β1 are in close proximity (<40 nm), the DNA oligonucleotides on the probes are ligated to form a circular DNA molecule. Perform rolling-circle amplification using a polymerase to amplify the circular DNA from the Duolink® In Situ Red Starter Kit. The amplified products were hybridized with fluorescently labeled oligonucleotides for visualization, then image and analyze the fluorescent signals using a confocal fluorescence microscope.

#### Statistics

Statistical analyses were performed using GraphPad software. Data were initially evaluated with the Shapiro-Wilk normality test and equal variance test. If both criteria were met, a two-tailed Student’s t-test was used for two-group comparisons. If either test failed, the Mann-Whitney rank-sum test was applied. For multiple-group comparisons, one-way ANOVA with Dunnett’s multiple comparisons was employed.

EC₅₀ values were calculated using GraphPad Prism software. For ATP assays, each individual plate or well was considered a biological replicate. For protein expression experiments, each sample or independent experiment was treated as a biological replicate. The statistical tests used, the number of biological replicates for each assay, and P-values for significant results are provided in the Figure Legends.

