## Supplementary material for "Screening of a kinase inhibitor library in human Huntington’s disease iPSC-derived striatal precursor neurons reveals a neuroprotective effect of PKC-α/β1 inhibition": All supplemental files

[1]Division of Neurobiology, Department of Psychiatry and Behavioral Sciences, Johns Hopkins University School of Medicine, Baltimore, Maryland 21287, USA.
[2]Department of Pharmacology and Molecular Sciences, Johns Hopkins University School of Medicine, Baltimore, Maryland 21287, USA.
[3]Department of Neuroscience, Johns Hopkins University School of Medicine, Baltimore, Maryland 21287, USA.
[4]Department of Neurology, Johns Hopkins University School of Medicine, Baltimore, Maryland 21287, USA.
[5]Department of Pathology, Johns Hopkins University School of Medicine, Baltimore, Maryland 21257, USA.
[6]These authors contributed equally.
[7]Lead contact.

**Supplementary Figure 1. HD ISPNs showed increased stress-induced apoptotic cell death than controls.**
ISPNs with 33/18 CAG repeats were seeded in 96-well plates, subjected to stress induction through growth factor and nutrient withdrawal (W/O) for 24 hours. Caspase 3/7 activities were quantified using the Caspase-Glo® 3/7 Assay System. Results are expressed as MEAN ± SEM and normalized to the 33Q under normal growth conditions (W), n=5-6. Statistical significance is analyzed by one-way ANOVA. $*p < 0.05$ vs. 180Q W.

**Supplementary Figure 2. GO6976 provided neuroprotection to HD human mature striatal neurons.**
ISPNs with 33/18 and 180/18 CAG repeats were differentiated for 14 days, subjected to stress induction through BDNF withdrawal (W/O), and treated with GO6976 at concentrations of 0.5 µM, or vehicle (Veh), for 24 hours. ATP levels were quantified using the CellTiter-Glo cell viability assay. Results are expressed as MEAN ± SEM and normalized to 33Q under normal differentiation conditions (Con), n = 5-6. One-way ANOVA, $*p < 0.05$ vs. 33Q W/O+Veh, $\#p < 0.05$ vs. 180Q Con+Veh, and $**p < 0.05$ vs. 180Q W/O +Veh.

**Supplementary Figure 3. GO6976 elevated ATP levels in 33Q ISPNS.**
ISPNs with 33/18 CAG repeats were seeded in 96-well plates, subjected to stress induction through growth factor and nutrient withdrawal (W/O), and treated with GO6976 at concentrations of 0.25, 0.5, or 1 µM, or vehicle (Veh), for 24 hours. ATP levels were quantified

using the CellTiter-Glo cell viability assay. Results are expressed as MEAN ± SEM and normalized to the normal growth conditions (Con), n = 5-6. Statistical significance is analyzed by one-way ANOVA. **p < 0.05 vs. 33Q W/O+Veh.

**Supplementary Figure 4. GO6976 did not increase ATP levels in the absence of cells.** 1 uM GO6976 or vehicle (Veh) were added to ISPN culture SCM2 medium for 24 hrs. ATP levels were quantified using the CellTiter-Glo cell viability assay. Results are expressed as MEAN ± SEM and normalized to the Veh group (n = 6). Student’s t-test.

#### Supplementary Table 1. Hits from the 1st round of screening that showed rescue effect in 180Q HD Striatal precursor neurons

| | Item Name | M.W. | Formula | Pathway | Target |
|---|---|---|---|---|---|
| 1 | Nu 6027 | 251.28 | C11H17N5O2 | Cell Cycle/Checkpoint | Cyclin-Dependent Kinases |
| 2 | PD168393 | 369.22 | C17H13BrN4O | JAK/STAT Signaling | EGFR |
| 3 | OSI-930 | 443.44 | C22H16F3N3O2S | Tyrosine Kinase | c-Kit |
| 4 | Lenvatinib (E7080) | 426.85 | C21H19ClN4O4 | Tyrosine Kinase | VEGFR |
| 5 | Quercetin dihydrate | 338.27 | C15H10O7.2H2O | PI3K/Akt/mTOR Signaling | PI3K |
| 6 | Brivanib (BMS-540215) | 370.38 | C19H19FN4O3 | Tyrosine Kinase | VEGFR |
| 7 | A 77-01 | 286.33 | C18H14N4 | TGF-β / Smad Signaling | TGF-βR1(ALK5) |
| 8 | LKB1 (AAK1 dual inhibitor) | 339.36 | C20H13N5O | Chromatin/Epigenetics | Pim |
| 9 | PHA-767491 | 213.24 | C12H11N3O | Cell Cycle/Checkpoint | Cdc7 |
| 10 | Ruxolitinib phosphate | 404.36 | C17H21N6O4P | Chromatin/Epigenetics | JAK |
| 11 | D4476 | 398.41 | C23H18N4O3 | Stem Cell | CK1 |
| 12 | BMS345541 hydrochloride | 291.78 | C14H18ClN5 | Immunology/Inflammation | IκB/IKK |
| 13 | XL388 | 455.5 | C23H22FN3O4S | PI3K/Akt/mTOR Signaling | mTOR |
| 14 | Go 6976 | 378.43 | C24H18N4O | TGF-β / Smad Signaling | PKC |
| 15 | BML-277 | 363.8 | C20H14ClN3O2 | Cell Cycle/Checkpoint | Chk |
| 16 | AZD1080 | 334.37 | C19H18N4O2 | PI3K/Akt/mTOR Signaling | GSK-3 |
| 17 | CC-401 | 388.47 | C22H24N6O | MAPK Signaling | JNK |
| 18 | AZD3759 | 459.9 | C22H23ClFN5O3 | JAK/STAT Signaling | EGFR |
| 19 | PS-1145 | 322.75 | C17H11ClN4O | Immunology/Inflammation | IκB/IKK |
| 20 | Decernotinib (VX-509) | 392.38 | C18H19F3N6O | Chromatin/Epigenetics | JAK |
| 21 | RG 13022 | 266.29 | C16H14N2O2 | JAK/STAT Signaling | EGFR |
| 22 | Kenpaullone | 327.18 | C16H11BrN2O | Cell Cycle/Checkpoint | Cyclin-Dependent Kinases |
| 23 | KPT-9274 | 610.62 | C35H29F3N4O3 | Cell Cycle/Checkpoint | PAK4 |

**Supplementary Figure 1.**
**HD ISPNs showed increased stress-induced apoptotic cell death than controls**

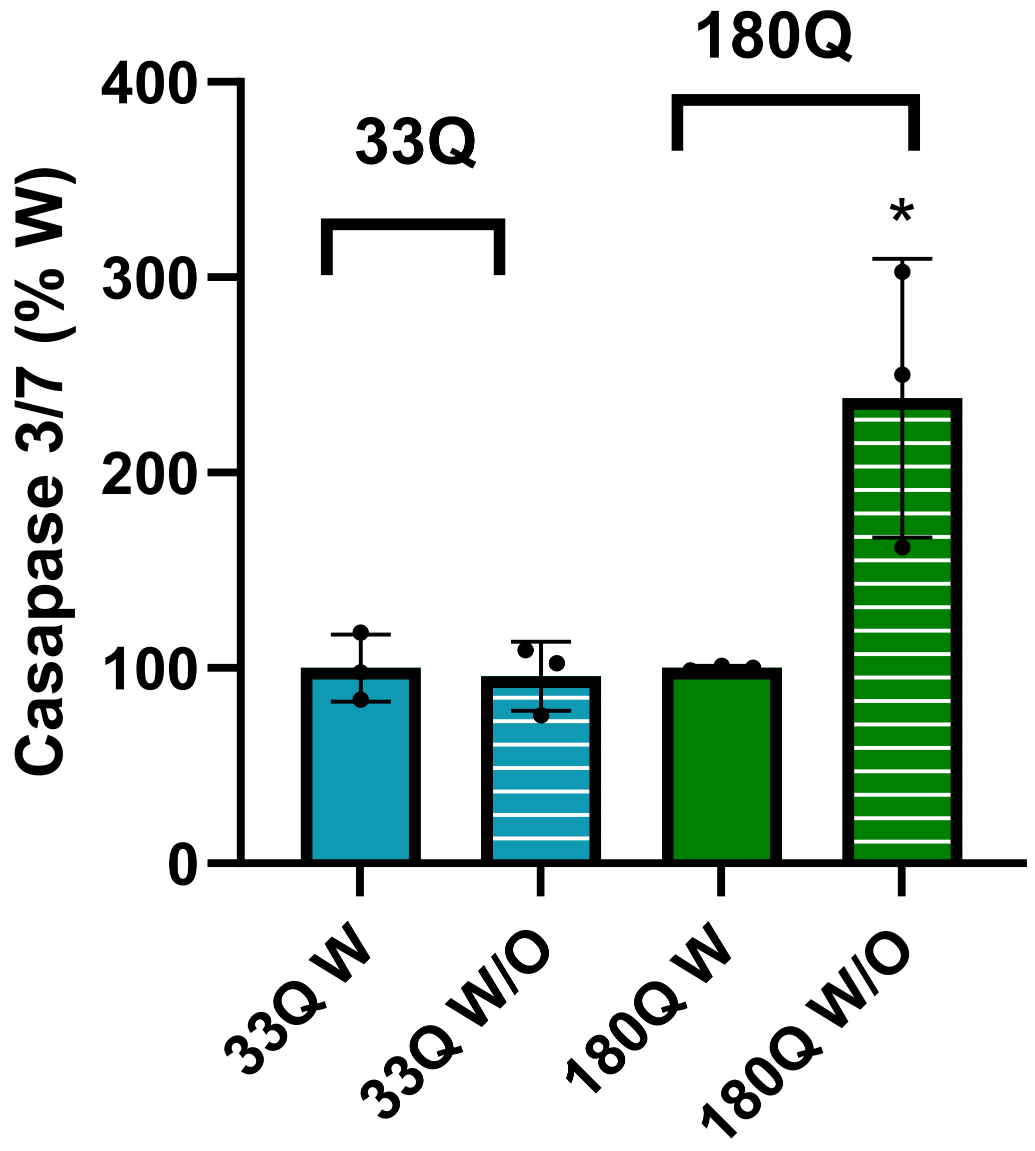

### Supplementary Figure 2.
### GO6976 provided neuroprotection to HD human mature striatal neurons

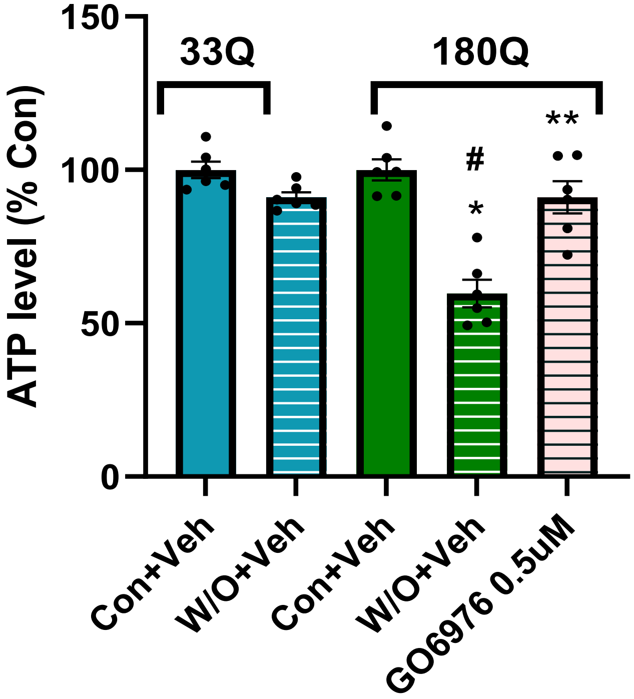

**Supplementary Figure 3.**
**GO6976 elevated ATP levels in 33Q ISPNS**

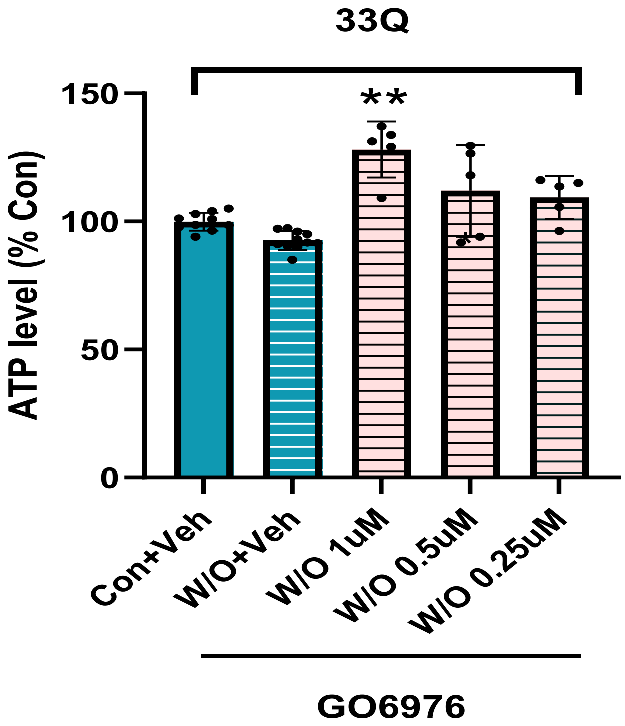

**Supplementary Figure 4.**
**GO6976 did not increase ATP levels in the absence of cells**

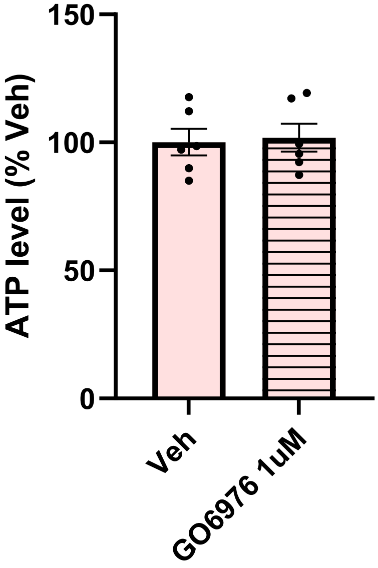
